# Loss-of-SIRT7 sensitizes hepatocellular carcinoma to sorafenib through the regulation of ERK Phosphorylation

**DOI:** 10.1101/2023.03.13.531998

**Authors:** Yuna Kim, Kwan-Young Jung, Yun Hak Kim, Pan Xu, Yunju Jo, Baeki E. Kang, Navin Pandit, Jeongho Kwon, Karim Gariani, Joanna Gariani, Junguee Lee, Jef Verbeek, Seungyoon Nam, Sung-Jin Bae, Ki-Tae Ha, Hyon-Seung Yi, Minho Shong, Kyun-Hwan Kim, Doyoun Kim, Chang-Woo Lee, Hee Jung Jung, Kwang Rok Kim, Kristina Schoonjans, Dongryeol Ryu, Johan Auwerx

## Abstract

The FDA-approved oral multi-kinase inhibitor, sorafenib (BAY 43-9006, Nexavar), is the first approved systemic therapy for patients with unresectable hepatocellular carcinoma (HCC). Although it has been shown to significantly improve the overall survival of patients with HCC, drug resistance limits the response rate to this therapeutic. Here, we report that acquired sorafenib resistance is associated with overexpression of the deacetylase, SIRT7, and a high level of ERK phosphorylation. Further, we identify that the hyperactivation of ERK is controlled by SIRT7-mediated deacetylation of DDX3X. The inhibition of SIRT7 combined with sorafenib resulted in a marked reduction of cell viability in vitro and of tumor growth in vivo. It seems plausible that SIRT7 is responsible for the acquired sorafenib resistance and its inhibition is most likely beneficial together in conjunction with sorafenib by suppressing ERK signaling.

**Highlights:**

- Sorafenib resistance in HCC is associated with SIRT7 and ERK hyperactivation.
- Suppression of SIRT7 combined with sorafenib restores sensitivity to sorafenib.
- SIRT7 controls sorafenib resistance through ERK activation by mediating DDX3X deacetylation.

## Introduction

Hepatocellular carcinoma (HCC) is the sixth most common malignancy and the fourth most common cause of cancer-related death in the world ^1^. The 5-year survival rate for advanced HCC is lower than 2%, which is ranked the lowest among all solid tumors ^2^. Unfortunately, advanced HCC is only eligible for systemic therapy which offers marginal survival benefit ^3^. Sorafenib is a multi-kinase inhibitor that mainly targets kinase activity in RAF/MEK/ERK signaling pathway and remains a globally accepted as the systemic first-line therapy for HCC for more than a decade. However, sorafenib monotherapy has shown to provide only modest improvement in overall median survival of about 2.8 months and its acquired resistance has become an obstacle to extending the overall survival benefits for HCC patients ^3,4^. Despite the effort to develop therapeutic strategies towards efficiently targeting chemoresistance of HCC, the heterogeneity of HCC limits uncovering driver genes and consequently, HCC remains one of the more challenging cancers to treat.

Meanwhile, findings from the Cancer Genome Atlas (TCGA) database suggest that SIRT7 expression is tightly correlated with the development of various types of cancer ^5–7^. Proteins of the mammalian sirtuin family (comprising SIRT1–SIRT7), known as NAD^+^-dependent deacetylases, are critical regulators implicated in a wide range of biological processes ^8^ and each of the sirtuins has shown considerably different functions and catalytic activities ^9^. SIRT7 is the least studied sirtuin family member and in human HCC, the expression of SIRT7 has been shown to be upregulated in a large cohort of HCC patients ^10,11^. Additionally, SIRT7 was reported to induce extracellular signaling-regulated kinase (ERK) phosphorylation and to activate the RAF/MEK/ERK signaling pathway to promote cancer cell proliferation ^12^.

The RAF/MEK/ERK pathway is heavily involved in tumor invasion and metastasis in HCC and phosphorylated ERK (pERK) is a well-known key downstream component of the RAF/MEK/ERK signaling cascade that reflects the level of pathway activation in tumor tissues. Phosphorylated ERK levels increased stepwise in accordance with metastatic potential of HCC cells ^13,14^ and accordingly, pERK levels were proposed to predict the responsiveness to sorafenib in HCC ^15^. However, this remains a contradictory issue in the clinic ^14,16–18^.

Nonetheless, little is known about the biological functions of SIRT7 in chemoresistance in HCC and its association with ERK activation in sorafenib-resistant HCC. Recently, a combination cancer therapy, using atezolizumab and bevacizumab, has been identified against treatment-resistant HCC that inhibit tumor growth and increases survival beyond what each individual component could have achieved independently ^19^ ^20^. In line with this, we examined the therapeutic potential of SIRT7 and evaluated whether it can be a key modulator in combination with sorafenib for mediating ERK activation in RAF/MEK/ERK signaling, influencing as such resistance mechanisms. We suggest that SIRT7 is a promising target in HCC and that its inhibition is relevant in the setting of sorafenib-resistant HCC.

## Results

### TCGA Liver Hepatocellular Carcinoma (TCGA-LIHC) data implies that SIRT7 expression may be associated with sorafenib resistance

Drug resistance still poses a significant obstacle to the long-term efficacy curative cancer treatments. The resistance to sorafenib, which is one of the first treatment options for HCC, has been reported through the last decade ^21^. Only a few studies have used the TCGA data to analyze the underpinning molecular candidates that could be the basis of sorafenib resistance. To candidates that potentially contribute to sorafenib resistant HCC, we analyzed TCGA-LIHC data then selected datasets based on sorafenib-treated patients who survived for more than 1000 days or died within 500 days and who might have sorafenib resistance (Figure 1A). Then, we conducted an unbiased gene set enrichment analysis (GSEA) to understand either sorafenib sensitive or resistance-associated gene sets. Among the 1,064 gene sets that belong to the biological process of gene ontology term (GO biological process), GSEA indicated that 235 gene sets were qualified statistically (Figure 1B). Interestingly, 75.1% of the gene sets among the 235 GSEA selected gene sets were associated with the biological function of the sirtuins ^8,22,23^, including gene sets of mitochondria (8.9%), epigenetics and transcription (11%), DNA repair and telomeres (9.3%), rRNA and ribosomes (7.7%), and cell cycle and replication (38.2%). The gene sets of macromolecule diacylation, mitochondrial gene expression, histone deacetylation, and ribosome biogenesis were enriched in short-term survivors, who might have sorafenib-resistance. Notably, all those selected gene sets were reported as downstream of SIRT7 ^24–29^. To see whether SIRT7 expression is associated with sorafenib resistance, we further analyzed the SIRT7 expression in three independent data sets (Figure 1D). While the expression of other protein deacetylases did not generate any associations, elevated SIRT7 expression was associated with sorafenib resistance (Figure 1E and S1A-S1C). Consistently, the TCGA-LIHC data indicate that SIRT7 expression has a negative correlation with the survival probability of patients treated with sorafenib (Figure 1F and G). This analysis suggests that SIRT7 may contribute, at least in part, to sorafenib resistance.

**Figure 1.**
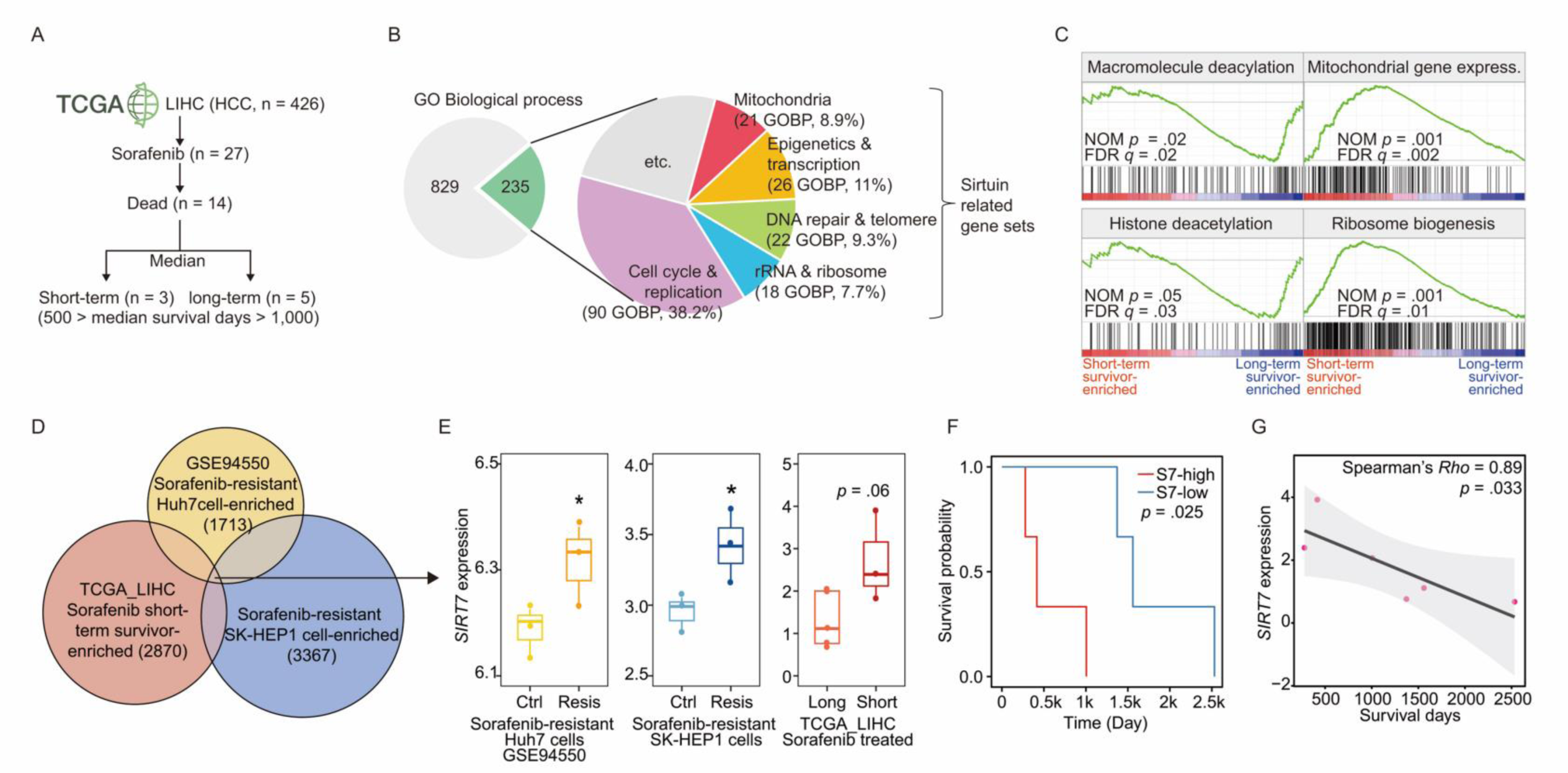
Association analysis of the transcriptional expressions in sorafenib resistant HCC samples. (A) Workflow summarizing the data selection process of the cancer genome atlas liver hepatocellular carcinoma (TCGA-LIHC). (B) Pie charts highlighting gene enrichment analysis (GSEA) selected gene sets (Gene ontology biological process term, GOBP). (C) Representative enrichment plots of GSEA that were enriched in short-term survivors who received sorafenib treatment in TCGA-LIHC data. (D and E) Venn diagram (D) showing RNA expression data of sorafenib short-term survivor enriched from TCGA-LIHC, sorafenib resistant Huh7 cells enriched RNA-seq data deposited in the Gene Expression Omnibus (GSE94550), and our RNA-seq data from sorafenib resistant SK-Hep1 cells and box plots (E) presenting SIRT7 expression. (F and G) Line graphs showing the survival probability (F) and the correlation (G) between hepatic SIRT7 and survival days.

### Acquired resistance to sorafenib in HCC cells is associated with SIRT7 expression

To investigate whether SIRT7 has potential to induce sorafenib resistance mechanism, we first tested the impact of alterations in SIRT7 expression in acquired sorafenib resistant HCC cells. By exposing Huh7 and SK-Hep1 HCC cell lines to increasing concentrations of sorafenib, we generated sorafenib-resistant HCC sub cell lines, named Huh7^SR^ and SK-Hep1^SR^ (Figure 2A). Attained sorafenib resistance was determined by the shift of the half maximal inhibitory concentration (IC_50_) toward a higher sorafenib concentration in all the resistant cell lines compared with their parental cell lines (Figures 2B and 2E). We then analyzed gene and protein expression of SIRT7 and established that SIRT7 was over-expressed in both Huh7^SR^ and SK-Hep1^SR^ cells in comparison with their parental cells (Figures 2C, 2D, 2F, and 2G). Together, these findings suggest that there is a negative association between SIRT7 expression and sorafenib responsiveness in HCC cells.

**Figure 2.**
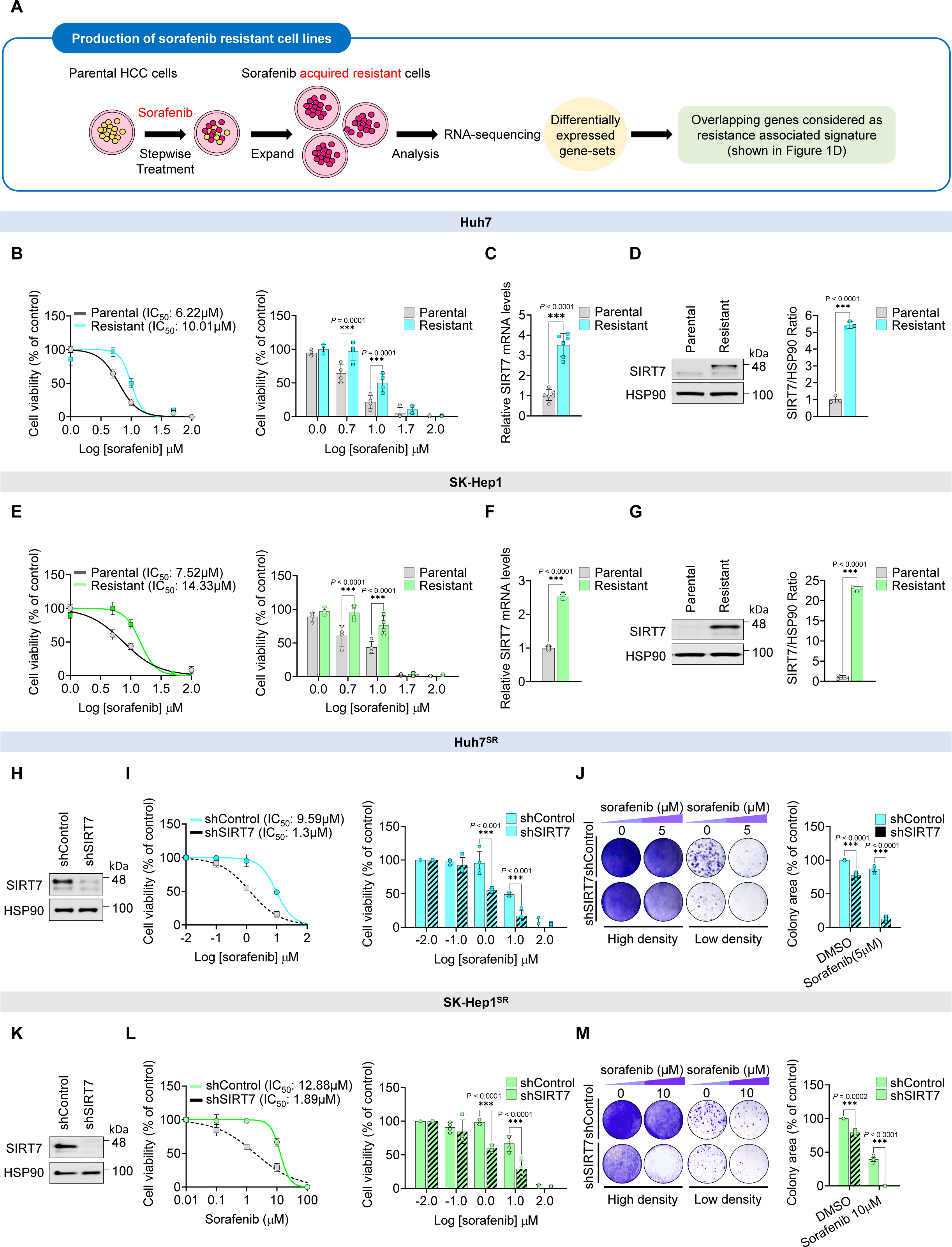
SIRT7 silencing inhibits proliferation of sorafenib resistant liver cancer cells. (A) Schematic image of the sorafenib adaptation process. Sorafenib parental HCC cell lines were exposed to sorafenib typically using a process of dose escalation. Overlapping genes considered as resistance associated signature were detected by means of RNA sequencing. (B and E) Huh7^SR^, SK-Hep1^SR^, and their parental cells were treated with increasing dose of sorafenib for 3 days. Relative cell viability was analyzed using MTT assay. The significance of the difference in IC50 values was determined. (C and F) The mRNA levels of SIRT7 in Huh7^SR^, SK-Hep1^SR^, and their parental cells were determined by RT-qPCR analysis. L32 which encodes *RPL32* was used as internal control. (D and G) The protein expression levels of SIRT7 in Huh7^SR^, SK-Hep1^SR^, and their parental cells were analyzed using western blotting. (H and K) The efficiency of SIRT7 knockdown in Huh7^SR^ and SK-Hep1^SR^ were determined by western blotting. (I and L) Comparison of sorafenib sensitivity in Huh7^SR^ and SK-Hep1^SR^ cells and SIRT7 knockdown Huh7^SR^ cells for 3 days. Relative cell viability was analyzed using MTT assay. (J and M) Huh7^SR^ and SK-Hep1^SR^ cells were cultured with or without sorafenib and synergistically combined with shRNA targeting SIRT7 for 10 days to assess viability by a colony formation assay. Data are mean ± SEM. Statistical analysis was conducted using the two-way ANOVA with post-hoc two-tailed *t*-test or two-tailed Student’s *t*-test where appropriate, \**P* < 0.05; \*\**P* < 0.01; \*\*\**P* < 0.001, compared with the control group.

### Inhibition of SIRT7 effectively re-sensitizes HCC to sorafenib

Based on the inverse association between SIRT7 expression and sorafenib response, we next questioned whether inhibition of SIRT7 can effectively restore the sorafenib sensitivity in acquired resistant HCC cells and whether it is synergistic with sorafenib. To address this question, both Huh7^SR^ and SK-Hep1^SR^ cells were transduced with a SIRT7 short hairpin RNA (shRNA) vector (Figures 2H and 2K) and were then cultured with or without sorafenib. Although SIRT7 knockdown alone could only offer modest reductions in cell viability and growth, suppression of SIRT7 expression in combination with sorafenib induced a marked inhibition of cell proliferation in the sorafenib unresponsive HCC cells in both short-term (Figures 2I and 2L) and long-term assays (Figures 2J and 2M). This, confirms that SIRT7 is responsible for sorafenib acquired resistance and that inhibition of SIRT7 has synergistic effects with a low dose of sorafenib effectively re-sensitizing HCC to sorafenib.

### SIRT7i combined with sorafenib drives a potent synergistic inhibition of cell viability

To test the therapeutic potential of SIRT7 inhibition on sorafenib resistant HCC cells, we produced two SIRT7 targeted chemical inhibitors (SIRT7i), named SIRT7i #1 and SIRT7i #2 (Figures 3A, 3C, S3A, and S3B). Although both SIRT7 inhibitors could attenuate cell viability when administered alone (Figures 3B and 3D), according to the results from colony formation assays (Figures 4A, 4D, 4G, and 4J), a potent synergistic inhibition of cell viability was seen when SIRT7i was combined with sorafenib (Figures 4B, 4E, 4H, and 4K). This effect was similar to what we observed upon knockdown of SIRT7 earlier. Further, the chemosensitivity in both sorafenib resistant HCC cells was also significantly increased by combination therapy measured by cell growth (Figures 3E, 3F, 3G, and 3H) and viability assays (Figures 4C, 4F, 4I, and 4L). Together, our data suggest that SIRT7i with sorafenib combination therapy is most likely to be effective in sorafenib resistant HCC.

**Figure 3.**
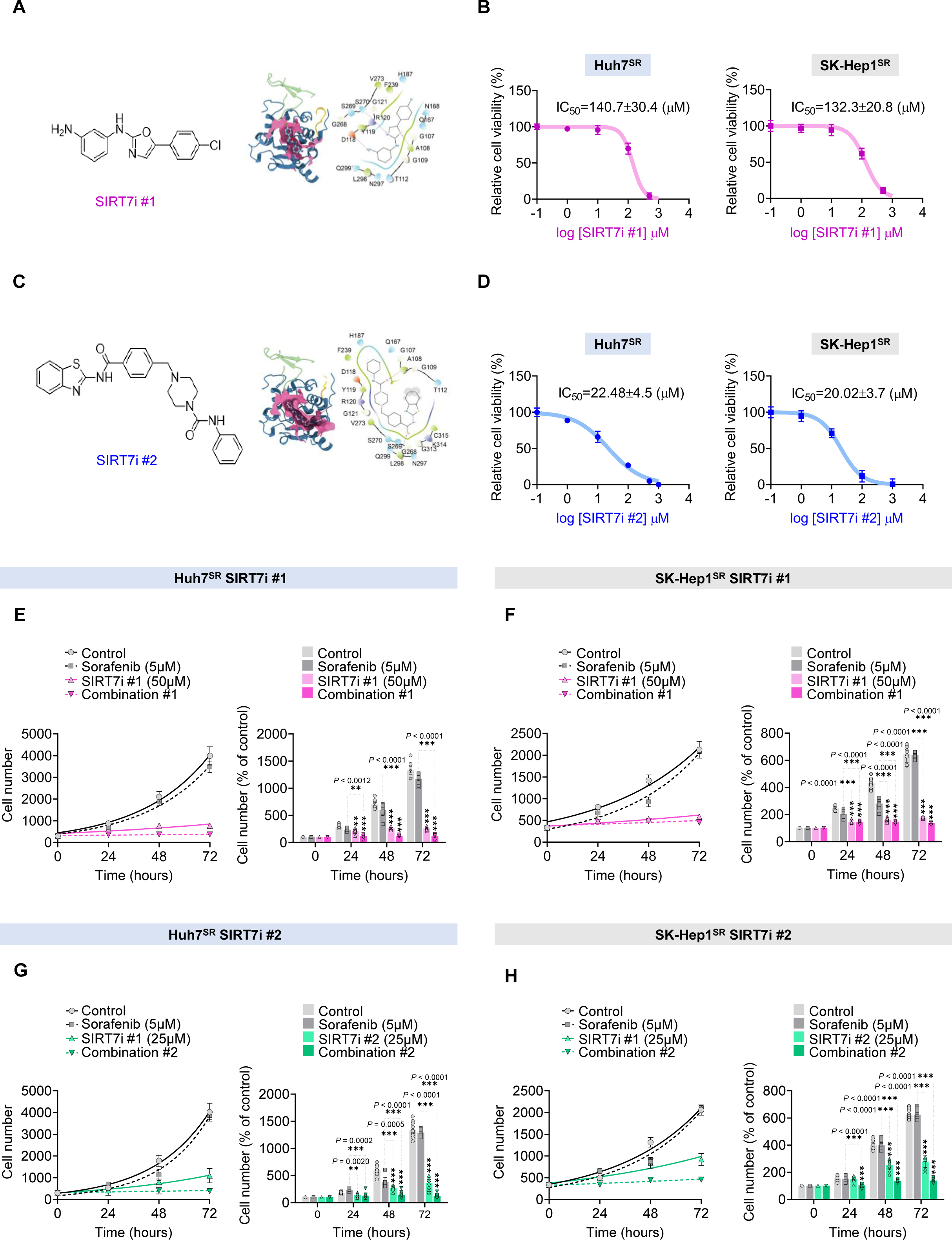
Development of SIRT7 targeted chemical inhibitors. (A and C) The chemical structure and docking pose with the interaction interface of SIRT7i #1 and SIRT7i #2. Theresidues involved in interaction between SIRT7i #1 and SIRT7 were represented by surface model and colored pink (Right). The 2D schematic representation of interaction interface between SIRT7i #1 or SIRT7i #2 and modeled SIRT7 structure. (B and D) Effect of SIRT7i #1 or SIRT7i #2 on cell viability. Both Huh7SR and SK-Hep1SR cells were treated with the indicated concentrations of SIRT7i #1 or SIRT7i #2 for 48h then cell viability was determined by MTT assay. (E and F) The cell numbers of Huh7SR and SK-Hep1SR treated with sorafenib or SIRT7i #1 alone and their combinations, indicated by Combinations #1, were determined by trypan blue exclusion assay at the indicated number of days thereafter. (G and H) The cell numbers of Huh7SR and SK-Hep1SR treated with sorafenib or SIRT7i #2 alone and their combinations, indicated by Combinations #2, were determined by trypan blue exclusion assay at the indicated number of days thereafter. Data are mean ± SEM. Statistical analysis was conducted using the two-way ANOVA with post-hoc two-tailed *t*-test, \**P* < 0.05; \*\**P* < 0.01; \*\*\**P* < 0.001, compared with the control group.

**Figure 4.**
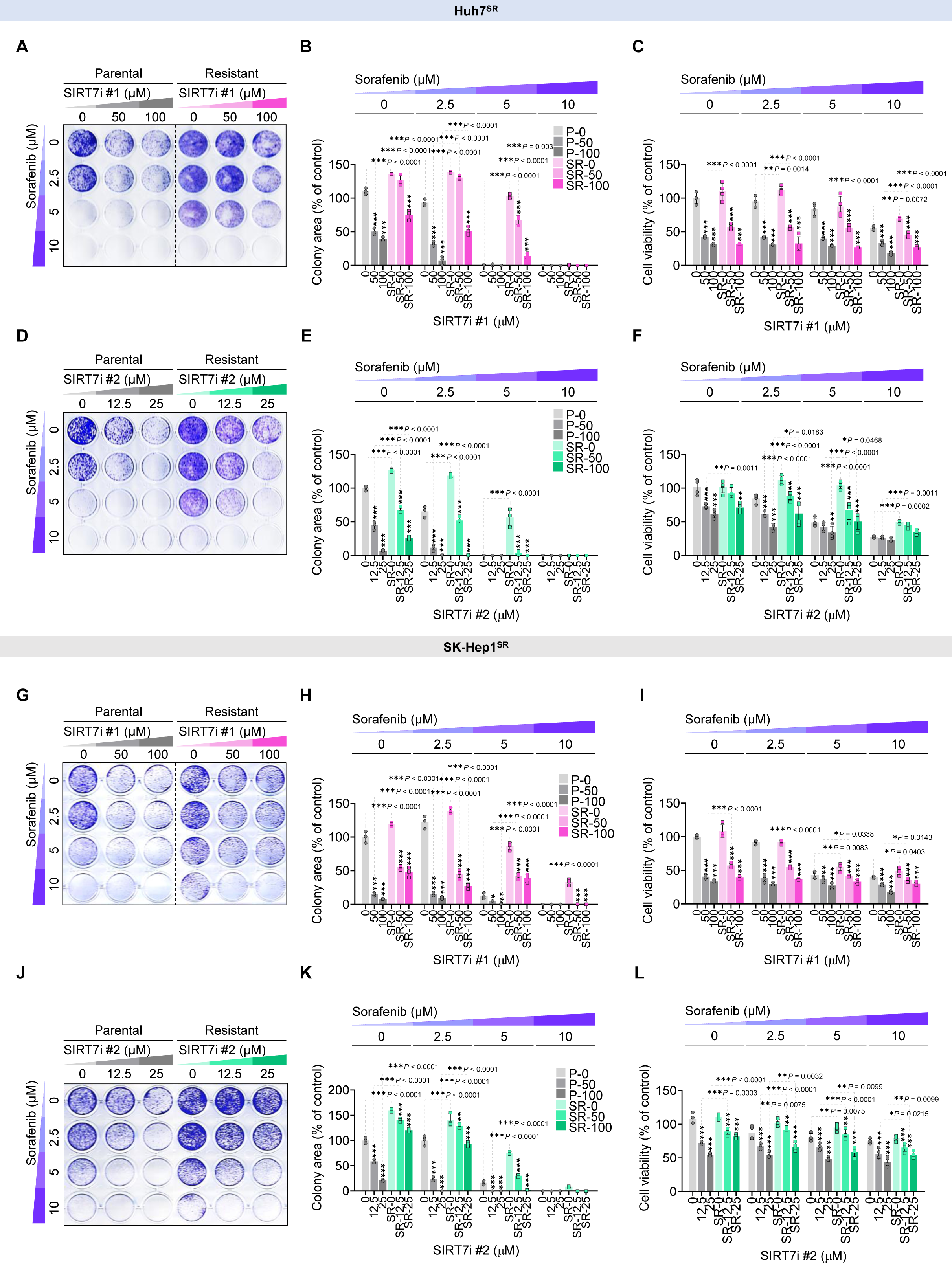
Chemical inhibitor of SIRT7 sensitizes sorafenib resistance. (A and D) Long-term colony formation assay of Huh7SR cells treated with sorafenib and indicated SIRT7 inhibitors. The cells were fixed and stained after 10-14 days. (B and E) The quantification of long-term colony formation assay shown in A and D. (C and F) Cell viability of Huh7SR treated with sorafenib and indicated SIRT7 inhibitors alone or their combinations at increasing concentrations were analyzed by MTT assay. (G and J) Long-term colony formation assay of SK-Hep1SR cells treated with sorafenib and indicated SIRT7 inhibitors. The cells were fixed and stained after 10-14 days. (H and K) The quantification of long-term colony formation assay shown in G and J. (I and L) Cell viability of SK-Hep1SR treated with sorafenib and indicated SIRT7 inhibitors alone or their combinations at increasing concentration were analyzed by MTT assay. Data are mean ± SEM. Statistical analysis was conducted using the two-way ANOVA with post-hoc two-tailed *t*-test, \**P* < 0.05; \*\**P* < 0.01; \*\*\**P* < 0.001, compared with the control group.

### SIRT7 contributes to sorafenib resistance through ERK activation

The level of ERK phosphorylation reflects the activation status of RAF/MEK/ERK signaling cascade, which is directly targeted by sorafenib. To investigate the predictive role of phosphorylated ERK levels on acquired resistance to sorafenib, we initially explored whether ERK hyperactivation is required for sorafenib resistance in HCC. We observed that hyperactivation of ERK corresponded with enhanced SIRT7 expression (Figures 5A and 5B) and importantly, decreased phosphorylation level of ERK was the consequence of inhibition of SIRT7 activity in sorafenib-resistant HCC cells compared to their parental cells (Figures 5C and 5D). To further understand whether overexpressed SIRT7 could be responsible for the poor response to sorafenib in HCC, the resistant HCC cells were treated with SIRT7i alone or combination of SIRT7i with sorafenib. Immunoblotting revealed that the inhibition of SIRT7 together with low dose of sorafenib treatment caused a more potent and complete inhibition of ERK phosphorylation compared to SIRT7i alone (Figures 5E and 5F). These data together suggest that SIRT7 could act as a driver for regulating ERK signaling in sorafenib acquired resistance.

**Figure 5.**
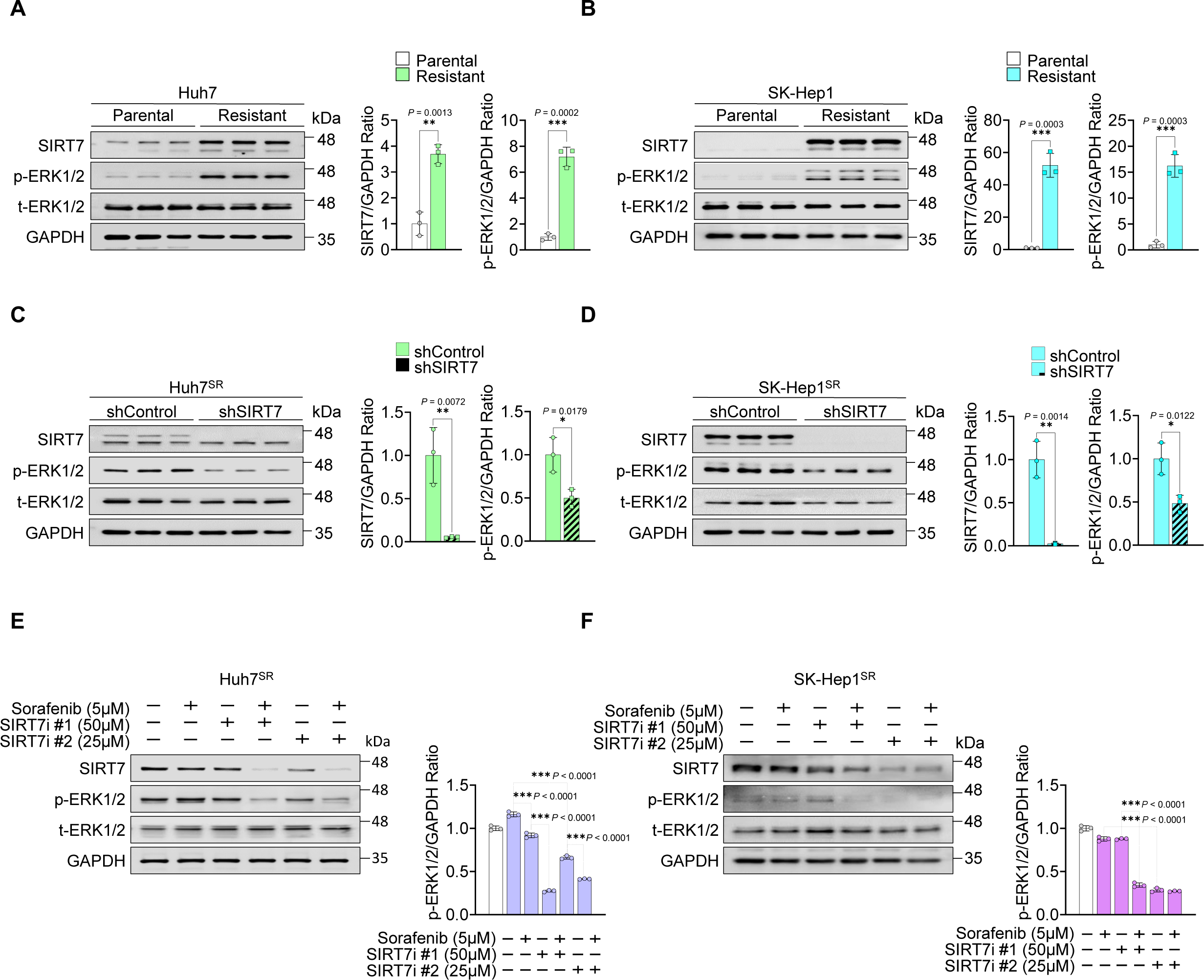
SIRT7 act as a key determinant of sorafenib resistance through pERK activation. (A) The expressions of both SIRT7 and pERK in Huh7SR cells were compared with their parental cells using western blotting. The levels of pERK were also quantified and normalized to GAPDH. (B) The expressions of both SIRT7 and pERK in SK-Hep1SR cells were compared with their parental cells using western blotting. The levels of pERK were also quantified and normalized to GAPDH. (C) Knockdown effect of SIRT7 in Huh7SR cells using the retrovirus and the expressions of pERK was analyzed by western blot. The levels of pERK were also quantified and normalized to GAPDH. (D) Knockdown effect of SIRT7 in SK-Hep1SR cells using the retrovirus and the expressions of pERK was analyzed by western blot. GAPDH served as a loading control. The levels of pERK were also quantified and normalized to GAPDH. (E) Huh7_SR_ cells were treated with sorafenib (5 μM), the SIRT7 inhibitor #1 (50 μM), the SIRT7 inhibitor #2 (25 μM) or their combination for 24 h, and western blot analysis was performed with the indicated antibodies. The levels of pERK were also quantified and normalized to GAPDH. (F) SK-Hep1SR cells were treated with sorafenib (5 μM), the SIRT7 inhibitor #1 (50 μM), the SIRT7 inhibitor #2 (25 μM) or their combination for 24 h, and western blot analysis was performed with the indicated antibodies. The levels of pERK were also quantified and normalized to GAPDH. Data are mean ± SEM. Statistical analysis was performed by two-tailed Student’s *t*-test, \**P* < 0.05; \*\**P* < 0.01; \*\*\**P* < 0.001, compared with the control group.

### Suppression of SIRT7 inhibits tumor growth through ERK phosphorylation

To address whether the in vitro findings can also be confirmed in vivo, Huh7^SR^ cells were injected into immunodeficient nude mice. Upon tumor establishment, xenografted mice were treated with control, sorafenib, SIRT7i #1, SIRT7i #2 or their combinations for 14 days (Figure 6A) and photographs of the mice, taken on day 14 were shown in Figure 6B. The combination of sorafenib with two SIRT7i showed a significant synergistic effect on tumor growth rate and tumor mass (Figures 6C and 6D) without affecting body weight (Figure 6H). To support these findings, immunohistochemical staining in each tumor tissues was also performed (Figure 6E). Whereas treatment of mice with SIRT7i alone showed moderate effects on proliferation and tumor progression, the combination treatments caused potent inhibition of tumor proliferation and progression, indicated by Ki-67 staining, a marker of proliferation and by AFP staining, a marker of tumor progression (Figures 6F and 6G). Likewise, immunoblot analysis also confirmed the strong suppressive effect of the combination treatments in tumor proliferation which resulted in robust reduction in ERK phosphorylation (Figures 6I and 6J).

**Figure 6.**
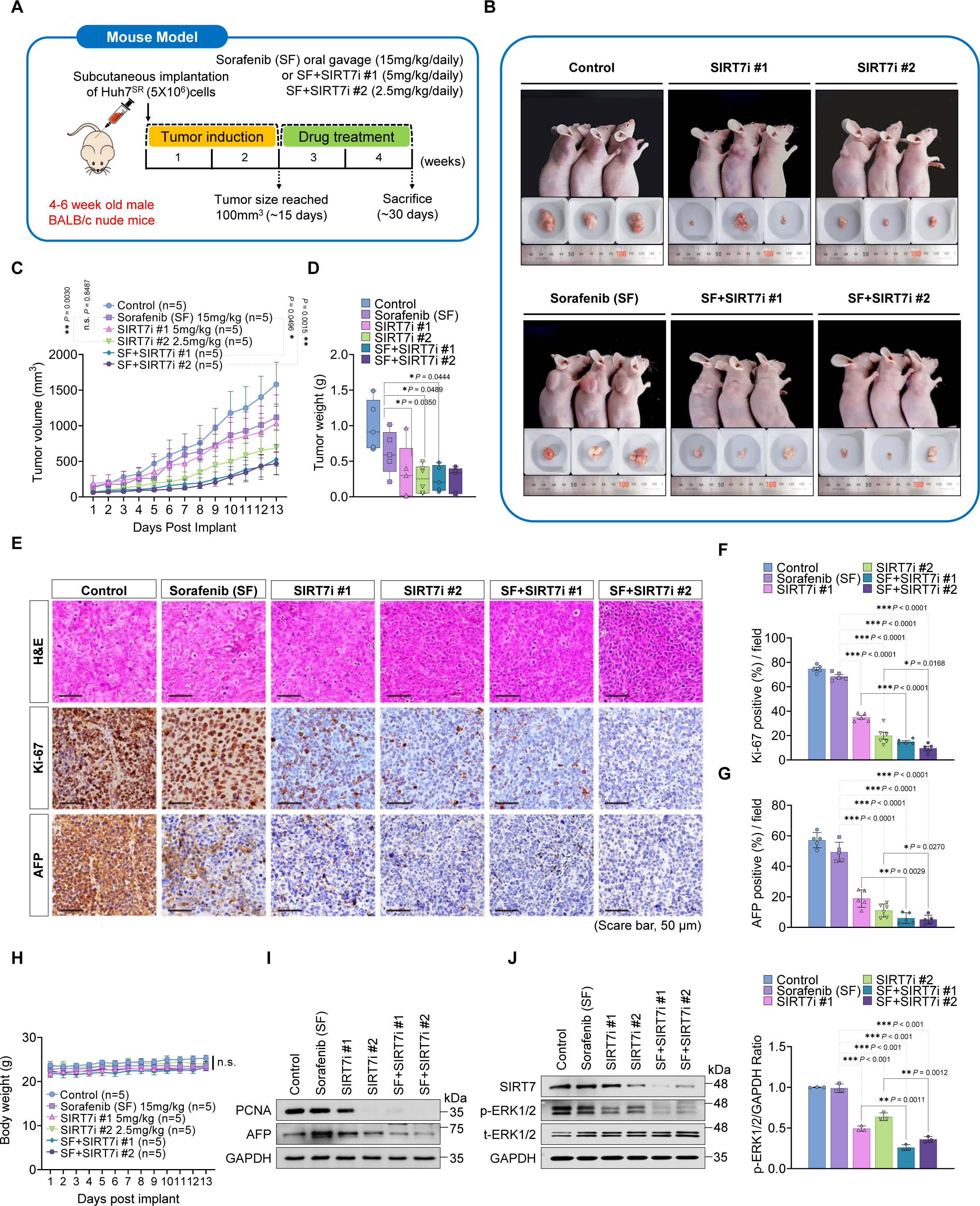
The combination of the SIRT7i and sorafenib displays potent anti-proliferative effects in xenograft mouse model. (A) Schematic image of Huh7^SR^ xenografted model. After tumor establishment (∼ 100 mm^3^), mice were treated with vehicle, sorafenib (15 mg kg^−1^), SIRT7i #1 (5 mg kg^−1^) or SIRT7i #2 (2.5 mg kg^−1^) plus sorafenib (15 mg kg^−1^) for about 13 days. n = 5-6 mice per group. (B) Representative photograph image of tumors formed in nude mice on day 14 for the experiment following injection of Huh7^SR^ cells in the right under armpit. (C) Growth curves of Huh7^SR^ xenografted mouse model. Xenografts were treated with control, sorafenib, SIRT7i #1, SIRT7i #2 or their combinations for about 13 days. (D) Huh7^SR^ xenografted tumors were excised and weighed. (E) Representative images of H&E, Ki-67, and AFP staining of tumors in the xenograft models. Scale bars, 50 μm. (F and G) Quantification of Ki-67 positive cells (F), AFP positive cells (G) per high-power field in representative sections from each group. n = 5 per group. (H) Body weights of mice in the xenografted mouse model with the aforementioned treatments were assessed. n = 5-6 mice per group. (I) Western blot analysis of PCAN and AFP levels in mouse tumor tissues. (J) Western blot analysis of total SIRT7 and pERK levels in mouse tumor tissues. The levels of pERK from each group were also quantified and normalized to GAPDH (Right). Data are mean ± SEM. Statistical analysis was conducted using the two-way ANOVA with post-hoc two-tailed *t*-test or two-tailed Student’s *t*-test where appropriate, \**P* < 0.05; \*\**P*< 0.01; \*\*\**P* < 0.001, compared with the control group.

### DDX3X is deacetylation target of SIRT7

Sirtuin enzymes are known to regulate in various metabolic processes through deacylation in target proteins ^8^. Therefore, the identification of their target substrates is a key for understanding sirtuin function. To further clarify the molecular mechanism underlying the synergistic effect of SIRT7 inhibition, we used interactome screening to identify a possible SIRT7 interactor that has an association with ERK signaling. According to the results from the interactome analysis based on the overlap between the acetylated and MARylated proteomes with the SIRT7 interactome (Figure 7A), DDX3X appeared to be a potent deacetylation candidate of SIRT7. We then validated DDX3X as a SIRT7 target using in vitro deacetylation assays by confirming that the p300-induced acetylation of DDX3X was reduced by SIRT7 (Figure 7A and 7B). As a direct binding partner of SIRT7 (Figure 7C), the level of DDX3X acetylation was analyzed by pulling down on the FLAG tag in FLAG-SIRT7 overexpressing HEK293 cells, verifying that the level of acetylated DDX3X was diminished upon overexpression of SIRT7 (Figure 7D). Conversely, knockdown of endogenous SIRT7 by transfecting an shSIRT7 vector resulted in increased levels of DDX3X pan-acetylation (Figure 7E), reconfirming that DDX3X is a direct deacetylation target of SIRT7. Through, mass spectrometric analysis we identified the acetylated site as lysine 55 in DDX3X (Figure 7F). We then produced the mutant of DDX3X to mimic the acetylated form by replacing lysine with glutamine (K55Q) (Figure 7G). As expected, enhanced overall acetylation was observed regardless of SIRT7 overexpression mediated deacetylation (Figure 7H).

**Figure 7.**
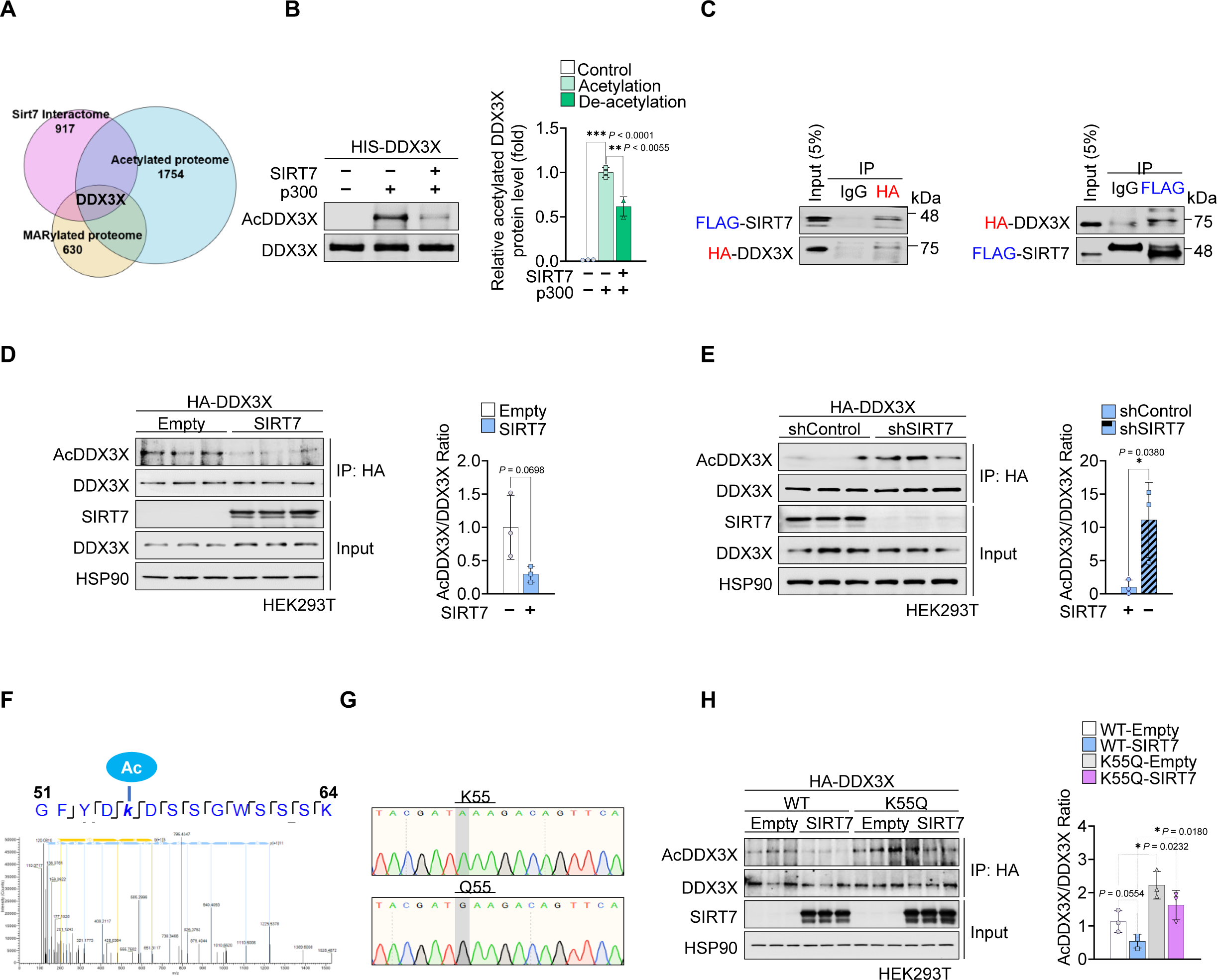
DDX3X is a deacetylation target of SIRT7. (A) Venn diagram showing the integration of acetylated proteome, MARylated proteome, and SIRT7 interactome. (B) In vitro deacetylation assay demonstrating that p300 acetylates and SIRT7 deacetylates DDX3X recombinant protein. (C) Identification of interaction between SIRT7 and DDX3X using immunoprecipitation (IP). HEK293T cells expressing FLAG-SIRT7 with HA-DDX3X were subjected to IP analysis. (D) Western blot analysis displaying reduced DDX3X acetylation in SIRT7 overexpressing HEK293T cells. (E) Western blot showing acetylated DDX3X in SIRT7 knockdown HEK293T cells. (F) Nano-LC-MS/MS analysis showing SIRT7-dependent in vitro deacetylation of AcK55 of DDX3X. (G) Genotype identification of the K55Q mutation (K55Q) in DDX3X plasmid. Electropherograms of sequencing displaying the point mutant site of DDX3X gene compared to wild-type (WT) sequencing. (H) Western blot analysis displaying expressions of acetylation status in WT or acetylation mimicking mutant K55Q of DDX3X upon SIRT7 overexpressing HEK293T cells. Data are mean ± SEM. Statistical analysis was performed by twotailed Student’s *t*-test, \**P* < 0.05; \*\**P* < 0.01; \*\*\**P* < 0.001, compared with the control group.

### SIRT7 mediates ERK signaling through DDX3X deacetylation

Previous studies have shown that DDX3X has a profound role in cancer cell invasion by regulating ERK signaling ^30^. Therefore, we hypothesized that decreased phosphorylated ERK levels due to SIRT7 inhibition may be linked with SIRT7-dependent DDX3X modification. To address our hypothesis, we initially tested whether SIRT7 regulated DDX3X protein stability. We transfected FLAG-SIRT7 or shSIRT7 into Huh7^SR^ and SK-Hep1^SR^ cells which were then treated with cycloheximide (CHX). The half-life of endogenous DDX3X was increased by SIRT7 overexpression (Figure 8A and 8C) whereas the knockdown of SIRT7 by shRNA reversed this effect (Figure 8B and 8D). We next investigated whether inhibition of SIRT7 mediated DDX3X acetylation is responsible for the regulation of ERK signaling upon sorafenib treatment. Upon transfection of Huh7^SR^ and SK-Hep1^SR^ with either DDX3X (WT) and DDX3X (K55Q) under sorafenib treatment, ERK phosphorylation was significantly more reduced in DDX3X (K55Q), indicating that K55 acetylation of DDX3X, mimicking SIRT7 deficiency, decreases ERK hyperactivation (Figures 8E and 8F).

**Figure. 8.**
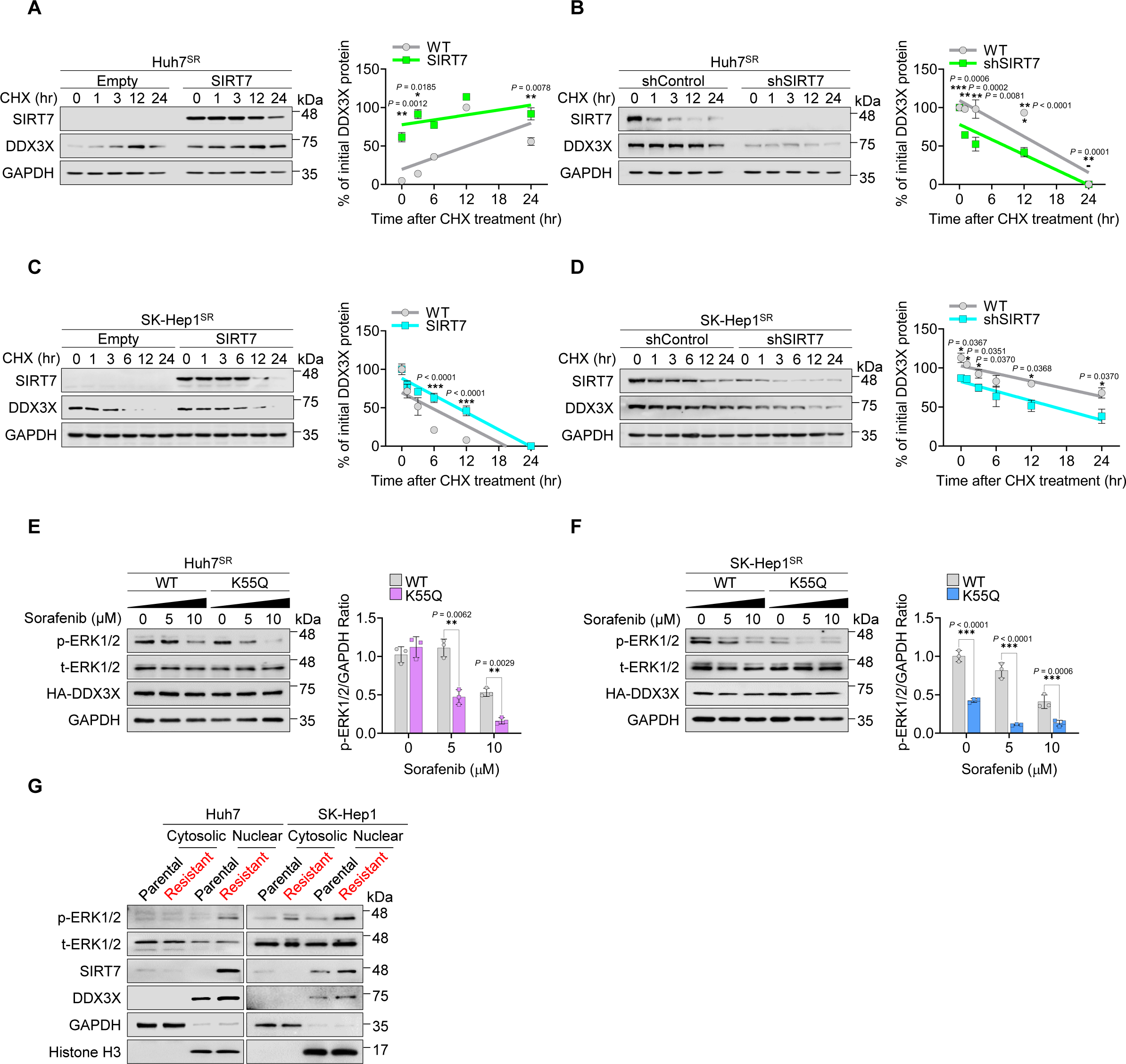
SIRT7 is required for activation of pERK signaling through stabilizing DDX3X. (A and C) Huh7^SR^ and SK-Hep1^SR^ cells were transfected with indicated plasmids and then treated with cycloheximide (CHX) (50 μg/mL) for the indicated times. Whole cell lysates were analyzed by western blotting. Quantitation of DDX3X protein levels was normalized to GAPDH. (B and D) Huh7^SR^ and SK-Hep1^SR^ cells were transduced with retrovirus mediated shRNA expressing SIRT7 and then treated with CHX (50 μg/mL) for the indicated times. Whole cell lysates were analyzed by western blotting. Quantitation of DDX3X protein levels was normalized to GAPDH. (E) Huh7^SR^ cells were transfected with WT or K55Q of DDX3X then treated with sorafenib for the indicated concentrations. Whole cell lysates were then analyzed by western blotting. Quantitation of pERK protein levels was normalized to GAPDH. (F) SK-Hep1^SR^ cells were transfected with WT or K55Q of DDX3X then treated with sorafenib for the indicated concentrations. Whole cell lysates were then analyzed by western blotting. Quantitation of pERK protein levels was normalized to GAPDH. (G) Comparison of Huh7^SR^ and SK-Hep1^SR^ with their parental cells in cytoplasmic and nuclear fractions which were subjected to immunoblotting as indicated. The data represent the means ± SEM. Statistical analysis was conducted using the two-way ANOVA with post-hoc two-tailed *t*-test or two-tailed Student’s *t*-test where appropriate, \**P* < 0.05; \*\**P*< 0.01; \*\*\**P* < 0.001, compared with the

Since SIRT7 is the only member of sirtuin family that is mainly localized in the nucleolus, we speculated that hyperactivated SIRT7 assembled with nuclear ERK (pERK) and DDX3X resulting in activation of the complex through protein-protein interactions. As we expected, pERK and DDX3X proteins were enriched in the nuclear fraction together with SIRT7 in sorafenib resistant cells compared to parental cells (Figure 8G). Also, a clear interaction between endogenous DDX3X and ERK proteins in Huh7^SR^ cells was demonstrated (Figure S5A). Overall, these findings support our hypothesis that inhibition of SIRT7 could repress hyperactive ERK signaling through acetylated DDX3X. Consequently, this restores sorafenib sensitivity and reduces the nuclear assembly of pERK and DDX3X required to maintain sorafenib resistance.

## Discussion

Here, we found that the sensitivity of acquired sorafenib resistance can be controlled by ERK signaling activated by SIRT7-mediated DDX3X deacetylation. The inhibition of SIRT7 activates an anti-survival signal in sorafenib-resistant HCC and is therefore synergistic with sorafenib.

### SIRT7 and ERK signaling

SIRT7 is possibly the least understood mammalian sirtuin, but its activity is important for human disease, particularly cancer. Increased SIRT7 expression is observed in many human cancers and a growing body of evidence suggests that it plays fundamental roles in oncogenic transformation and tumor biology ^10,31–33^. In HCC, SIRT7 expression is reported to be upregulated in a large cohort of patients with HCC, and increased SIRT7 is associated with increased tumor grade ^10^. Consistent with these findings, Zhao et al. also observed that both mRNA and protein levels of SIRT7 were significantly upregulated in most HCC tissues compared with adjacent nontumoral liver tissues ^11^. More importantly, SIRT7 expression is significantly correlated with poor overall survival in liver cancer cases ^11^. Our results show that SIRT7 expression is further enhanced in acquired sorafenib resistant HCC cells and this finding supports the earlier conclusions that SIRT7 acts as an oncogene in HCC development and its induction may be associated with hyperactivation of oncogenic signaling in drug resistant HCC.

The level of ERK phosphorylation indicates the activation status of the serine/threonine kinase RAF/MEK/ERK signaling cascade in liver cancer favoring HCC progression by promoting cellular proliferation, survival, tumor growth, and eventually resistance ^34^, which is directly targeted by sorafenib ^35^. However, controversial results were reported on the association between high basal ERK phosphorylation and survival benefits in patients treated with sorafenib ^17,18^. It has been shown that inhibition of cell proliferation upon sorafenib treatment is directly linked to the basal levels of pERK expression in patient-derived HCC cells indicating that sorafenib sensitivity could be dependent on the activation of the RAF/MEK/ERK signaling pathway measured by the basal pERK levels in HCC tumor cells ^14^. However, according to a recent sorafenib combination study, there was no strong correlation between the baseline pERK levels and sorafenib sensitivity observed; rather, pERK could only be a biomarker for assessing responsiveness of sorafenib in sorafenib combination therapy ^36^.

In line with the recent finding of pERK as an independent biomarker for sorafenib sensitivity, our data indicate that hyperactivation of both ERK and enhanced SIRT7 expression were seen in sorafenib acquired resistant HCC cells and ERK activity was well suppressed by a combination therapy of sorafenib with SIRT7 inhibitors, which leads to a strong synergistic lethal effect on resistant HCC cells. To date, no activating modulator of ERK has been described in sorafenib-resistant HCC; our new data ascertain the role of SIRT7 as pro-oncogenic regulator of ERK activation.

### SIRT7 dependent DDX3X regulation and ERK signaling

The involvement of DDX3X in activating the RAF/MEK/ERK signaling pathways to promote cancer invasion capability was previously examined in KRAS-mutated colon cancer cells ^30^ by demonstrating the increase of invasion capability in DDX3X-overexpressing cancer cells which was reversed by ERK inhibitor treatment or shERK transfection. In the present study, we identify additional evidence that overexpression of the DDX3X acetylation mimic K55Q protein in sorafenib-resistant HCC cells was sufficient to reduce ERK phosphorylation under sorafenib stimulation. This indicates that the decisive role of SIRT7 in modulating ERK activation involves epigenetic modulation of DDX3X.

Interestingly, it has been also shown that changes in cellular localization of DDX3X in normal tissues might lead to tumorigenesis ^37,38^ and subcellular localization of DDX3X to organelles, including the nucleolus, centrosome and mitochondria endows it with different functions based on its subcellular location ^39–41^. Remarkably, an early study found that DDX3X localized specifically to the nucleolus was linked to worse cancer prognosis ^40^. The nucleolus is also increasingly recognized as a target for cancer therapy ^42^ and many of essential proteins localized in the nucleolus are enriched for RNA processing, including DDX3X ^43^. The presence of high nuclear DDX3X levels may reflect increased protein synthesis demands in cancers ^40^, and SIRT7, the direct interacting partner of DDX3X, is also mostly enriched in nucleoli ^44^, which are increased in size and number in aggressive tumors ^42^. In our study clearly showed that DDX3X expression was restricted solely to the nuclear fraction, in accordance with hyperactivation of nuclear ERK and SIRT7 (Figure 8G). However, the exact role of DDX3X in the nuclear compartment and how the subcellular localization of DDX3X is regulated in tumorigenesis through modulating ERK signaling remains still to be elucidated. Further research is hence required to characterize the function of DDX3X related to subcellular compartmentalization and the cellular mechanisms behind increased nuclear retention of ERK which may facilitate the development of improved cancer strategies.

In summary, our data suggest that SIRT7 acts as a key effector molecule that regulates ERK activation by mediating DDX3X deacetylation. Suppression of SIRT7 activity combined with sorafenib could effectively restore sorafenib sensitivity. Thus, the combination therapy identified here may represent a promising strategy for treating advanced HCC associated with sorafenib resistance.

## Methods

### Cell Lines

The human HCC cell lines, Huh7 and SK-Hep1 were purchased from Korean Cell Line Bank (KCLB, Korea), human embryonic kidney cell line, HEK293, was obtained from ATCC. Cell lines were cultured in DMEM with 10% FBS and 1% penicillin/streptomycin in humidified incubator of 5% CO_2_ at 37°C. Originally, Sorafenib resistant cells, Huh7^SR^ and SK-Hep1^SR^ were kindly gifted from Prof. Keun-Gyu Park (Kyungpook National University, Korea) and these sublines were further generated by gradient exposures of sorafenib for about 6 months to acquire its resistance.

### Mouse Models

All mice were treated according to protocols reviewed and approved by the Institutional Animal Care and Use Committee (IACUC) of Sungkyunkwan University School of Medicine (SUSM) (code/SKKUIACUC 2021-07-47-1, approval date 20 Aug 2021). SUSM is an Association for Assessment and Accreditation of Laboratory Animal Care International (No. 001004) accredited facility. All experimental procedures were carried out in accordance with the regulations of the IACUC guidelines at Sungkyunkwan University. BALB/c nude mice (5-6-week-old, male, Charles River) were used for xenograft experiment.

### Cell proliferation assay

Sorafenib-resistant cells and their parental cells were seeded at a density of 5000 cells per well in 96 well plates. Briefly, equal number of cells were treated with various concentrations of sorafenib or SIRT7i #1 ^45^ and SIRT7i #2, which was given by Dr. Kwan-Young Jung, alone or their combinations. Cell viability was determined by MTT assay after 3 days of incubation and cell numbers were calculated by trypan blue exclusion assay at the indicated number of days thereafter.

### Colony formation assay

Cells were seeded into 12-well plates (10,000 cells per well) and cultured in the presence of drugs, as indicated. Then, the cell colonies were washed with PBS, fixed by ratio of acetic acid to methanol 1:3, and stained with crystal violet (Sigma).

### Plasmids and shRNA Retroviral transduction

Oligonucleotides were synthesized and cloned into the pSIREN-RetroQ retroviral shRNA expression vector (Clontech). Then pSIREN-RetroQ-Sirt7 and the negative control vector (pSIREN-RetroQ) were transfected into HEK293 cells using PEI transfection reagent (Sigma). The pSIREN-RetroQ-Sirt7 and the negative control vector (pSIREN-RetroQ) were gifted from Prof. Kazuya Yamagata (Kumamoto University, Japan) ^46^. Subsequently, sorafenib resistant HCC cells were infected with retroviral supernatants using 8 μg/ml polybrene. After 24 h of incubation, the supernatant was replaced by medium containing 2 μg/ml puromycin. After 48 h, selection of viral transduced cell lines was completed.

### Mass spectrometry analysis

Gel lanes were cut into pieces and subjected to in-gel digestion with endoproteinase. Peptide digests were resuspended and analyzed by nano LC-MS/MS with an Orbitrap Elite Mass Spectrometer (Thermo Fischer Scientific) coupled to an ultraperformance LC system (Thermo Fischer Scientific Ultimate 3000 RSLC). Data analysis was performed with Proteome Discoverer (v. 1.3).

### Site directed mutagenesis

The DDX3X expression plasmid, which was cloned by PCR into pcDNA3.1(+) vector using HindIII and XhoI restriction enzyme sites, was kindly donated by Dr. Kyun-Hwan Kim (Sungkyunkwan University, Korea) ^47^. The Muta-Direct™ Site Directed Mutagenesis Kit (INtRON, Korea) was used to generate DDX3X site-directed mutant according to the manufacturer’s instructions. Briefly, PCR was performed to create mutagenesis alongside with wide type (WT) of DDX3X plasmid as its template. Then, mutant and WT of DDX3X plasmids were transformed into XL-10GOLD cells. All the plasmids were confirmed by chromatogram DNA sequencing.

### In Vitro Acetylation and Deacetylation Assays

In vitro acetylation and deacetylation assays were performed as originally described ^48,49^. In brief, 1 mg of recombinant DDX3X protein was incubated with 500 ng of recombinant p300 in acetylation buffer (50 mM Tris-HCl [pH 8], 100 mM NaCl, 10% glycerol, 1 mM phenylmethylsulfonyl fluoride [PMSF], 1 mM dithiothreitol [DTT], 1 mg/ml bepstatin, 1 mg/ml leupeptin, 1 mg/ml pepstatin, 1 mM sodium butyrate, and 150 mM acetyl-CoA) for 1 hr at 37C. After incubation, samples were resolved on SDS-PAGE and analyzed by western blot or used for in vitro deacetylation assays. For deacetylation assays, 1 mg of acetylated DDX3X was incubated with 500 ng recombinant of SIRT7 protein in the deacetylation buffer (50 mM Tris-HCl [pH 9], 4 mM MgCl2, 0.2 mM DTT, 1 mg/ml bepstatin, 1 mg/ml leupeptin, 1 mg/ml pepstatin, and 1 mM NAD^+^) for 30 min with constant agitation. The incubated samples were resolved on SDS-PAGE and analyzed by western blot or used for mass spectrometry analysis. Recombinant proteins p300 and SIRT7 were kindly provided by Michael O. Hottiger (University of Zurich).

### Immunoprecipitation assay

Cells were harvested in immunoprecipitation buffer (20 mM HEPES pH 7.9, 200 mM NaCl, 0.5 mM EDTA, 10% glycerol, 0.2% NP-40) containing phosphatase and protease inhibitors (Sigma), and lysed for 30 min at 4°C. Lysate was cleared by two sequential centrifugation steps. HA-coupled magnetic beads (Thermo Fisher Scientific) or Protein A/G (sc-2003, Santa Cruz) were used and incubated overnight at 4°C on rotating wheel. The pull-down was eluted from beads in 2x Laemmli sample buffer (20 mM Tris pH 6.8, 4% SDS, 0.02% bromophenol blue, 13.4% glycerol, 2% β-mercaptoethanol) at 95°C for 5 min. Samples were analyzed by immunoblotting.

### Protein lysate preparation and western blotting

Cells were washed three times with PBS and lysed with RIPA buffer (20mM Tris-HCl pH7.5, 150mM NaCl, 1mM EDTA, 1% NP-40, 1% sodium deoxycholate, 1mM Na_3_VO_4_, 1mM NaF) supplemented with Complete Protease Inhibitor (Roche), Phosphatase Inhibitor Cocktails II and III (Sigma). Protein concentration in supernatants were quantified using *DC* protein assay kit (BioRad). Equal amounts of total proteins (20μg) were separated by SDS-PAGE and transferred to a 0.45-mm Immobilon-P PVDF Membrane (Millipore, Merck) and the membrane was incubated with primary antibodies overnight at 4°C. After incubation with HRP-conjugated α-rabbit or α-mouse secondary antibodies for 1 h at room temperature, target protein bands were detected using chemiluminescent detection Reagent for 5 min and visualized using imaging system (FUSION Solo S, VILBER).

### Isolation of RNA and qRT-PCR

Total RNA was extracted using Trizol Reagent (Invitrogen) and was reverse-transcribed into complementary DNA with random hexamer primer (Thermo Fisher Scientific). qRT-PCR was performed in triplicates using QuantStudio 6 Flex Real-Time PCR System (Applied Biosystems) according to the manufacturer’s instructions.

### RNA sequencing

Total RNA was prepared using the Trizol method and verified the size of PCR enriched fragments, together with the template size distribution by running on an Agilent Technologies 2100 Bioanalyzer using a DNA 1000 chip. (Agilent Technologies, Santa Clara, CA, USA). We performed next-generation sequencing using a HiSeq system according to the manufacturer’s instructions (Illumina, San Diego, CA, USA). Sequences were processed and analyzed by Macrogen (Seoul, Korea).

### Mouse xenografts

Huh7^SR^ cell (5×10^6^ cells per mouse) were injected subcutaneously into the right posterior flanks of 5-week-old BALB/c nude mice (male, 5-6 mice per group). Tumor volume based on digital caliper measurements was calculated using the modified ellipsoidal formula: tumor volume = ½ length × width^2^. After tumor establishment, mice were randomly assigned to 5 days per week treatment with vehicle, sorafenib (15 mg kg^−1^, oral gavage), SIRT7i #1 (5 mg kg^−1^, oral gavage), and SIRT7i #2 (2.5 mg kg^−1^, oral gavage) or a drug combination in which each compound was administered at the same dose and schedule as the single agent. Sorafenib and two SIRT7 inhibitors were dissolved in ethanol and Cremophor EL/95% (50:50) to prepare a stock solution (4×) and was diluted in sterile water to obtain the final concentration prior to use.

### Immunohistochemical staining

Tumor tissues were fixed, embedded, and sliced into 5 μm thick sections. Briefly, paraffin sections were first deparaffinized and then hydrated. After microwave antigen retrieval, endogenous peroxidase activity was blocked with incubation of the slides in 0.3% H_2_O_2_, and non-specific binding sites were blocked with 10% goat serum. Then the sections were incubated with primary antibodies (mouse anti-Ki-67and mouse anti-AFP) overnight at 4°C, followed by HRP-conjugated secondary antibodies for 1 h at room temperature. Finally, the sections were developed in diaminobenzidine solution under a microscope and counterstained with hematoxylin. The proliferation index was determined as number of Ki-67 or AFP-positive cells/total number of cells in 5 randomly selected high-power fields (magnification × 200) using ImageJ (v.1.46r) software.

### Statistical Analysis

For analyses involving multiple comparisons, one-way ANOVA with Bonferroni post-hoc test was used. Student’s t-test was used. Bar graphs show mean ± SEM, unless otherwise stated in the figure legend. All graphs were plotted and analyzed with GraphPad Prism 8 Software. p values <0.05 were considered statistically significant.

## Data and code availability

The RNA seq and TCGA data discussed in this publication have been deposited in NCBI’s Gene Expression Omnibus (GEO: GSE94550).

## Acknowledgements

This study was supported by the Basic Science Research Program through the National Research Foundation of Korea (NRF) funded by the Ministry of Science and ICT (NRF-2020R1A2C2010964 to D.R. and 2021R1A6A3A13044725 to Y.K.), the Ecole Polytechnique Federale de Lausanne (EPFL), the European Research Council (ERC-AdG-787702), the Swiss National Science Foundation (SNSF 31003A_179435), a GRL grant of the National Research Foundation of Korea (NRF 2017K1A1A2013124), and the Korea Research Institute of Chemical Technology (KRICT) funded by the Ministry of Science and ICT (BSF21-301 to K.R).

## Author contributions

Y.K., K.-Y.J., Y.H.K., K.R.K., K.S., J.A and D.R. conceived and designed the project. Y.K, K.-Y.J., Y.H.K., N.-K.L., S.W., P.X. and S.-J.B. performed experiments. Y.H.K, Y.J., B.E.K., J.A. and D.R. performed bioinformatic analysis. K.-Y.J., D.K. H.J.J. and K.R.K developed the SIRT7 specific inhibitors and tested *in vitro*. Y.K., Y.H.K., N.-K.L., S.W., P.X., K.G., S.-J.B., K.-T.H., K.-H.K., J.U.L., H.-S.Y., M.S., contributed for animal experiments, histological and phenotypic analyses. Y.K., K.-Y.J., Y.H.K., J.V., K.R.K., K.S., D.R., and J.A wrote and edited the manuscript with input from all other authors.

