## Supplementary Figures 1-5 for "Loss-of-SIRT7 sensitizes hepatocellular carcinoma to sorafenib through the regulation of ERK Phosphorylation"

**Supplementary Figure 1.**

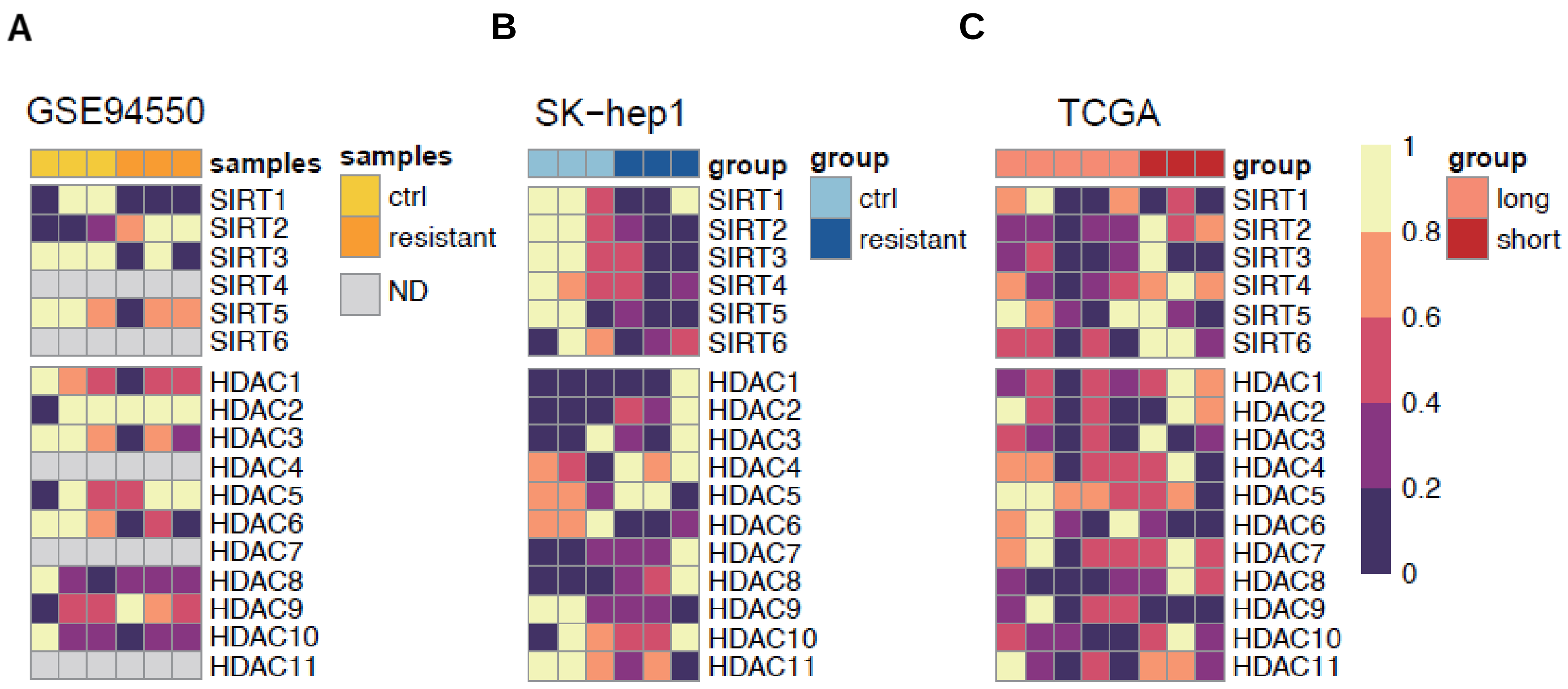


**Figure S1, related to Figure 1. The expression profiles of the protein deacetylase family.**

(A-C) Heatmaps showing the expression patterns of protein deacetylases in sorafenib-resistant Huh7 cells enriched RNA-seq data deposited in the Gene Expression Omnibus (GSE94550) (A), our RNA-seq data from sorafenib resistant SK-Hep1 cells (B), and in the livers of the sorafenib short-term survivors (C).

**Supplementary Figure 2.**

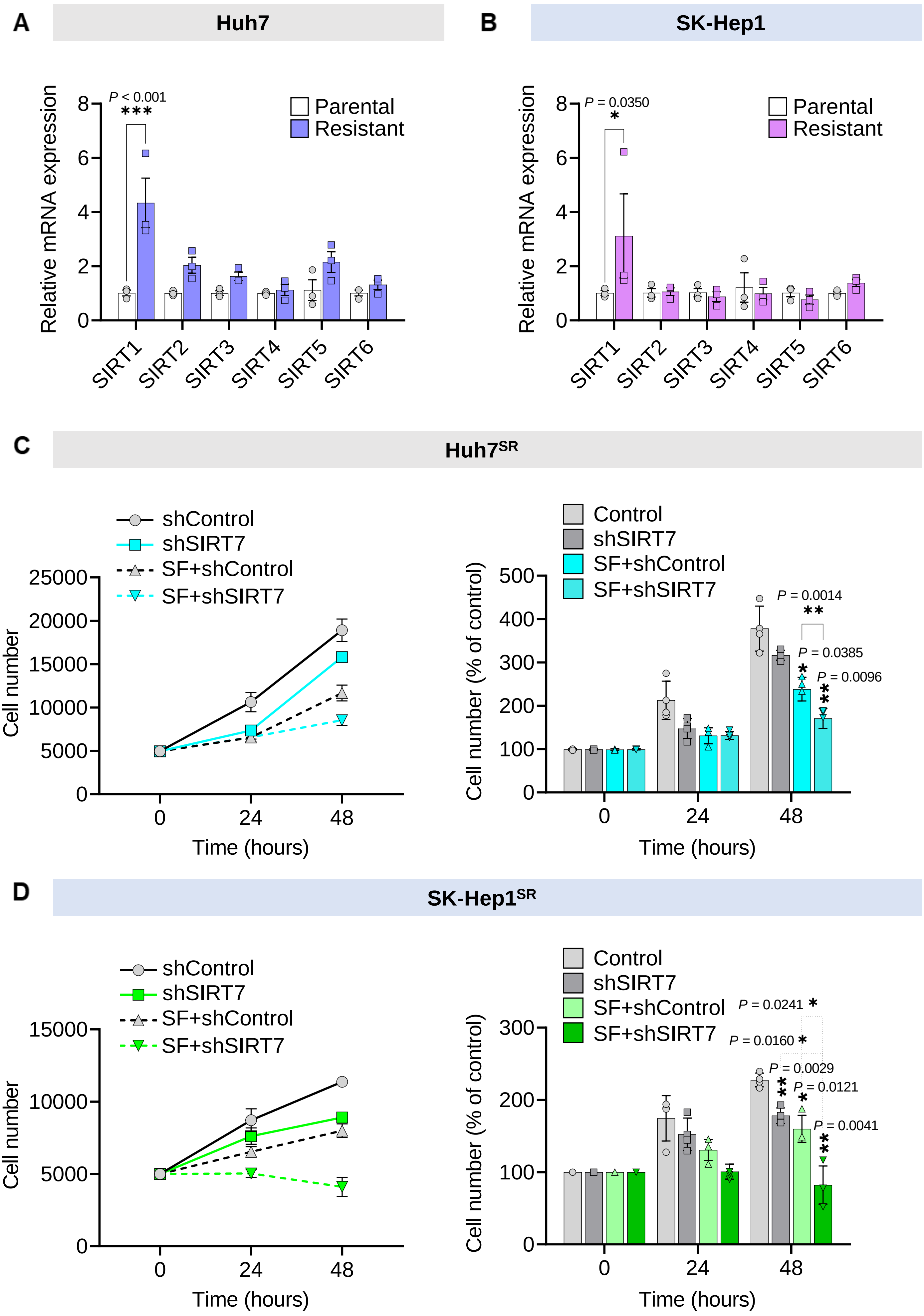

**Figure S2, related to Figure 2. SIRT7 silencing inhibits proliferation of sorafenib resistant liver cancer cells.**

(A) The mRNA levels of SIRT1-6 in Huh7 and Huh7$^{SR}$ cells were determined by RT-qPCR analysis. L32 which encodes *RPL32* was used as internal control.

(B) The mRNA levels of SIRT1-6 in SK-Hep1 and SK-Hep1$^{SR}$ cells were determined by RT-qPCR analysis. L32 which encodes *RPL32* was used as internal control.

(C) The cell numbers of Huh7$^{SR}$ treated with sorafenib or shSIRT7 alone and their combination were determined by trypan blue exclusion assay at the indicated number of days thereafter.

(D) The cell numbers of SK-Hep1$^{SR}$ treated with sorafenib or shSIRT7 alone and their combination were determined by trypan blue exclusion assay at the indicated number of days thereafter. Data are mean ± SEM. Statistical analysis was conducted using the two-way ANOVA with post-hoc two-tailed *t*-test or two-tailed Student's *t*-test where appropriate, *$P < 0.05$; **$P < 0.01$; ***$P < 0.001$, compared with the control group.

**Supplementary Figure 3.**

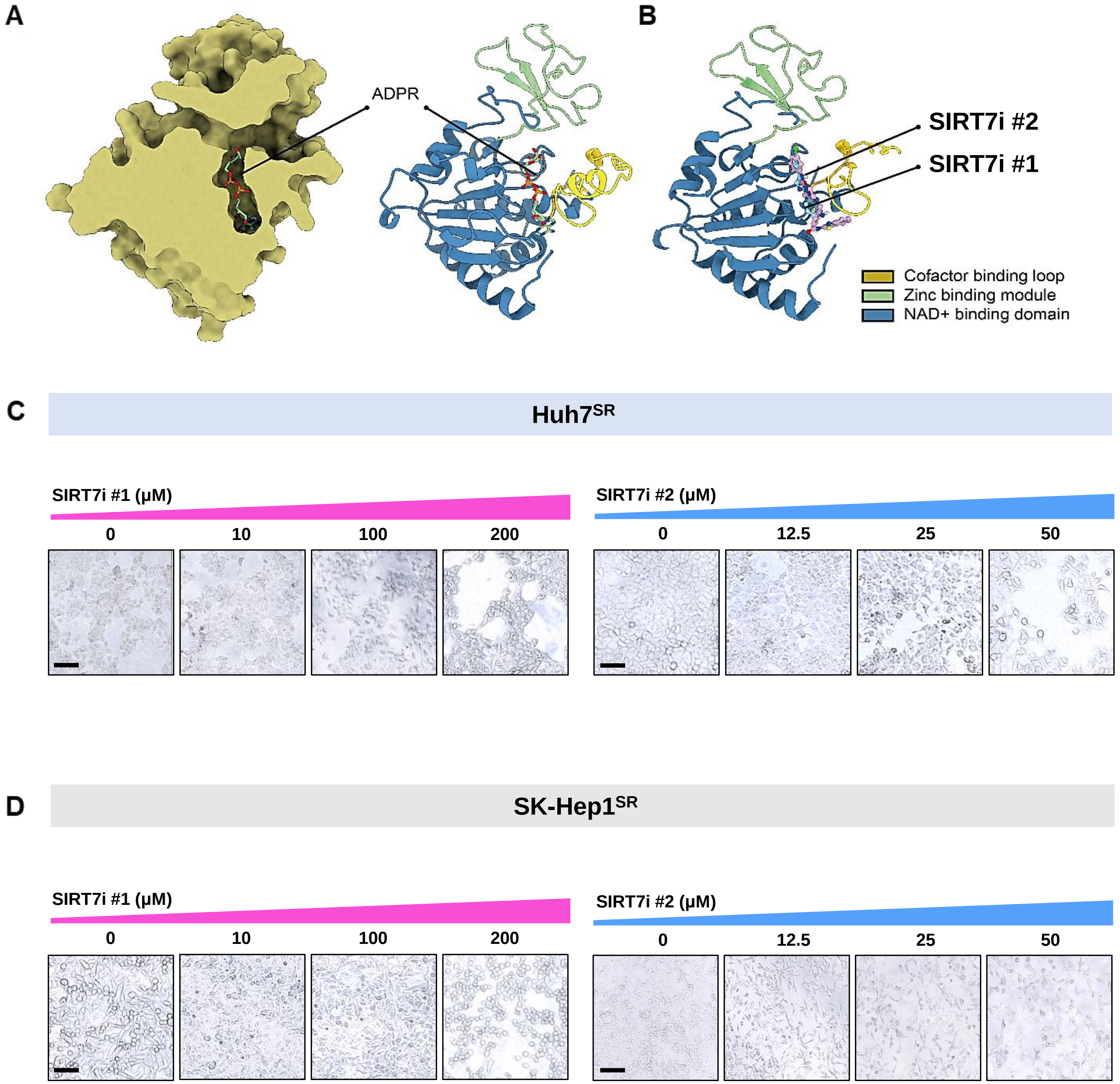

**Figure S3, related to Figure 3. Development of SIRT7 targeted chemical inhibitors.**

(A) The modeled structure of SIRT7 in complex with ADPR with surface (left) and ribbon (right) representation. The ADPR were labeled and represented by stick and ball model in the binding pocket of SIRT7 structure.

(B) The predicted binding poses of SIRT7i #1 and SIRT7 #2 in ADPR binding pocket of modeled SIRT7 structure. The SIRT7i #1 and SIRT7 #2 were represented by ball and stick model and colored by cyan and pink, respectively.

(C) Under light microscope with 10X magnification showing Huh7$^{SR}$ cells treated with either SIRT7i #1 or SIRT7 #2.

(D) Under light microscope with 10X magnification showing SK-Hep1$^{SR}$ cells treated with either SIRT7i #1 or SIRT7 #2.

### Supplementary Figure 4.

A

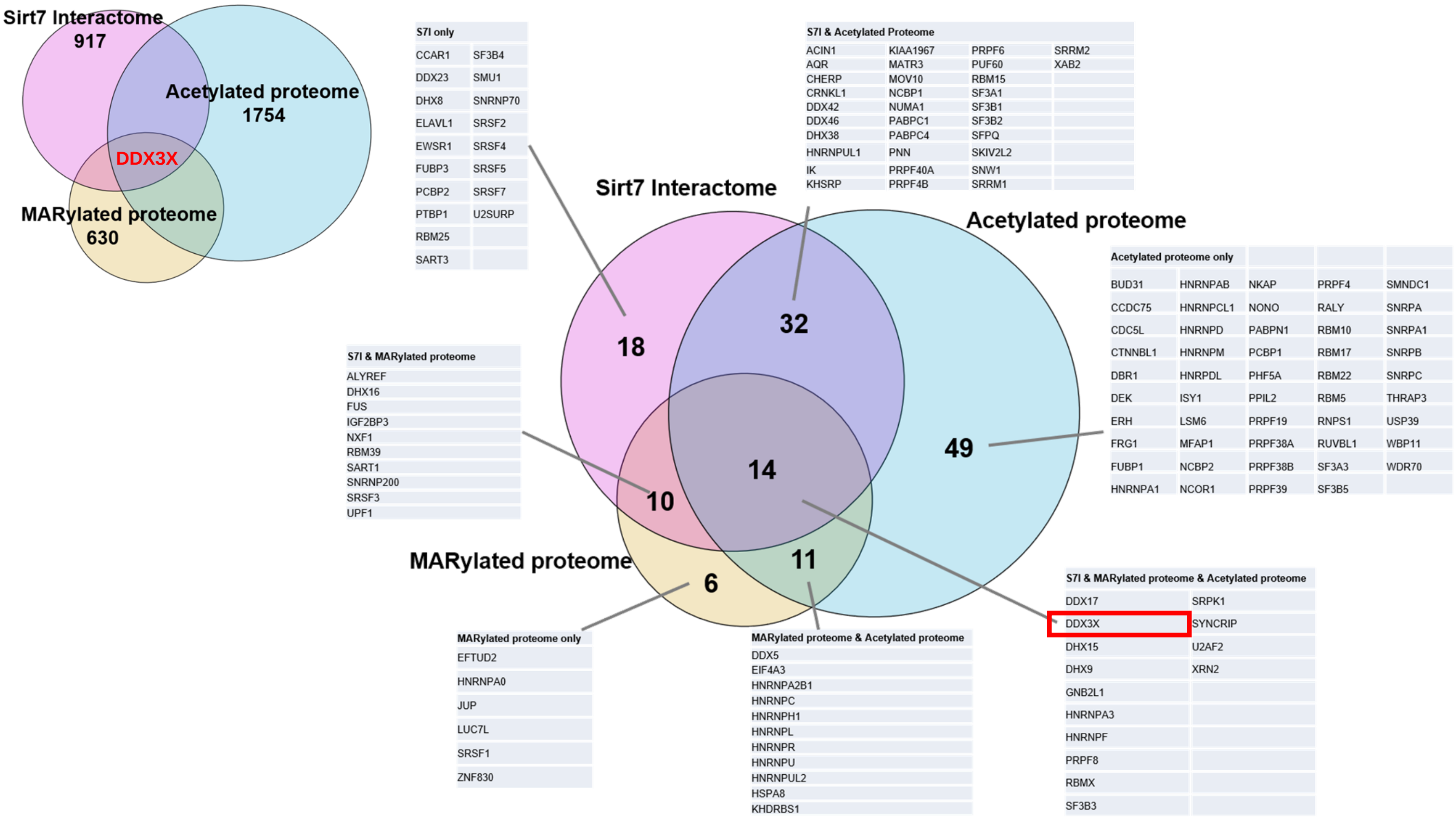


**Figure S4, related to Figure 7. DDX3X is deacetylation target of SIRT7**

(A) Acetylated proteome is based on previous large-scale mass spectrometry (MS) analysis. MARylated proteome is based on ADPRiboDB. SIRT7 interactome is constructed on BioGRID4.0 interaction datasets.

**Supplementary Figure 5.**

A

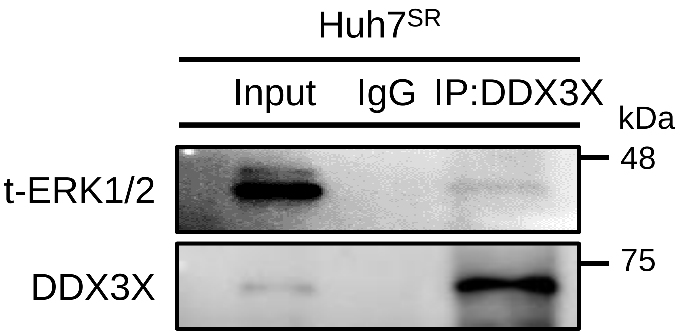


**Figure S5, related to Figure 8. SIRT7 is required for activation of pERK signaling through stabilizing DDX3X**

(A) Identification of interaction between DDX3X and ERK using IP. Co-IP of endogenous DDX3X with ERK in Huh7$^{SR}$ cells.
